# PKA and HOG signaling contribute separable roles to anaerobic xylose fermentation in yeast engineered for biofuel production

**DOI:** 10.1101/540534

**Authors:** Ellen R. Wagner, Kevin S. Myers, Nicholas M. Riley, Joshua J. Coon, Audrey P. Gasch

**Author notes:** These authors contributed equally to this work (Ellen R. Wagner, Kevin S. Myers).

## Abstract

Lignocellulosic biomass offers a sustainable source for biofuel production that does not compete with food-based cropping systems. Importantly, two critical bottlenecks prevent economic adoption: many industrially relevant microorganisms cannot ferment pentose sugars prevalent in lignocellulosic medium, leaving a significant amount of carbon unutilized. Furthermore, chemical biomass pretreatment required to release fermentable sugars generates a variety of toxins, which inhibit microbial growth and metabolism, specifically limiting pentose utilization in engineered strains. Here we dissected genetic determinants of anaerobic xylose fermentation and stress tolerance in chemically pretreated corn stover biomass, called hydrolysate. We previously revealed that loss-of-function mutations in the stress-responsive MAP kinase *HOG1* and negative regulator of the RAS/Protein Kinase A (PKA) pathway, *IRA2*, enhances anaerobic xylose fermentation. However, these mutations likely reduce cells’ ability to tolerate the toxins present in lignocellulosic hydrolysate, making the strain especially vulnerable to it. We tested the contributions of Hog1 and PKA signaling via IRA2 or PKA negative regulatory subunit BCY1 to metabolism, growth, and stress tolerance in corn stover hydrolysate and laboratory medium with mixed sugars. We found mutations causing upregulated PKA activity increase growth rate and glucose consumption in various media but do not have a specific impact on xylose fermentation. In contrast, mutation of *HOG1* specifically increased xylose usage. We hypothesized improving stress tolerance would enhance the rate of xylose consumption in hydrolysate. Surprisingly, increasing stress tolerance did not augment xylose fermentation in lignocellulosic medium in this strain background, suggesting other mechanisms besides cellular stress limit this strain’s ability for anaerobic xylose fermentation in hydrolysate.

## Introduction

Lignocellulosic biomass offers a sustainable source for bioenergy. The use of leftover agriculture byproducts and plants grown on marginal lands for biofuel production reduces waste and removes dependency on food-based cropping systems. Notably, there are two major bottlenecks for sustainable biofuel production from lignocellulosic material. First, many microbes, including industrially relevant *Saccharomyces cerevisiae*, cannot innately ferment pentose sugars like xylose, which comprise a significant fraction of the sugars released from deconstructed biomass (1). Second, the harsh chemical treatment of plant biomass required to release lignocellulosic sugars produces a variety of toxins and stresses that inhibit microbial growth and fermentation (2,3). A goal for the biofuel industry is to engineer stress-tolerant microbes to convert all available sugars to the desired products by routing cellular resources toward product formation and away from cell growth and other unnecessary physiological responses (4).

Several labs have evolved or engineered yeast for anaerobic xylose usage, introducing either xylose isomerase (5–7) or xylose reductase (XR) with xylitol dehydrogenase (XDH) (8–13), and over-expressing xylulokinase for increased flux (14–17). However, most strains do not use xylose unless further evolved through laboratory selection (7,18–21). In some cases, the genetic basis for evolved improvements in anaerobic utilization are known from sequencing of evolved lines. We previously engineered a stress-tolerant strain with XR and XDH (strain Y22-3) and evolved it for aerobic xylose respiration, producing strain Y127. To enable anaerobic xylose fermentation, Y127 was propagated anaerobically on xylose, generating strain Y128 (21). Aerobic xylose respiration in the Y127 strain was enabled by null mutations of the Fe-S cluster protein *ISU1* and the osmotic stress response MAP kinase *HOG1*. Maximal anaerobic fermentation in the evolved Y128 strain was facilitated by these mutations plus additional null mutations of the negative regulator of RAS/PKA signaling, *IRA2*, and *GRE3*, an aldose reductase that siphons xylose to xylitol (22). A subsequent study independently generated an anaerobic-xylose fermenting strain, confirmed the requirement of the *ISU1* deletion, and further found deletion of the upstream HOG pathway regulator *SSK2* improved xylose fermentation (23). Thus, mutations in these pathways play a generalizable role in anaerobic xylose fermentation across labs and strains.

While mutations that promote xylose utilization are known, the specific roles for each mutation and how the RAS/PKA and HOG pathways intersect to enable anaerobic xylose utilization remain unclear. RAS signaling promotes growth on preferred nutrients like glucose, in part by activating adenylate cyclase to produce cAMP, which binds to the PKA negative regulatory subunit Bcy1 to enable PKA activity (24). Ira1/2 are the GTPase activating proteins (GAPs) that inhibit Ras1/2 by converting GTP (RAS-active state) to GDP (RAS-inactive state). On the other hand, Hog1 is best characterized as an osmotic stress response MAP kinase and leads to the upregulation of stress-responsive transcription factors and other enzymes and defense systems (25). How Hog1 contributes to xylose fermentation is unknown, although the kinase was recently shown to play a role in the response to glucose levels (26–30). PKA and Hog1 have opposing roles on the stress response: PKA activates transcription factors required for growth-promoting genes and directly suppresses stress-activated transcription factors like Msn2/Msn4, while Hog1 activity induces stress-defense regulators and contributes to the repression of growth-promoting genes (31).

Increased stress sensitivity is a major limitation for industrial use of evolved strains with RAS/PKA and HOG mutations and a barrier to sustainable lignocellulosic bioenergy production. Chemical pretreatment of plant biomass is required to release fermentable sugars into the resulting “hydrolysate.” This treatment produces a variety of toxins and stressors that limit microorganisms’ ability to ferment, particularly impacting fermentation and growth during xylose consumption (2,32,33). One group of toxins are lignocellulosic hydrolysate inhibitors (or lignotoxins), which are released from breakdown of hemicellulose and cellulose and include furans, phenolics, and aliphatic acids. Lignotoxins disrupt central carbon metabolism pathways by generating reactive oxygen species and depleting the cells of ATP, NADH, and NADPH, in part through increased activity of ATP-dependent efflux pumps and detoxification (34–36), ultimately decreasing available resources for growth and metabolism. Therefore, strains must be tolerant to the toxins present in hydrolysate for efficient fermentation of lignocellulosic material, but the mutations required for efficient anaerobic xylose fermentation produce stress-susceptible strains. Upregulated PKA activity suppresses stress-defense pathways (37–40), while *HOG1* deletion decreases cells’ ability to mount a stress response. This likely has a direct impact on existing strains, limiting their industrial use. One possible solution is increasing stress tolerance in these strains will enable better anaerobic xylose fermentation in industrially relevant hydrolysates. Other groups have found varying levels of success improving toxin tolerance (3) through overexpression of stress response transcription factors (41,42), mitochondrial NADH-cytochrome b5 reductase (42), an oxidative stress protein kinase (43), and furaldehyde reductases (44), as well as through mating, gene shuffling, and evolution (45–48).

We recently discovered an alternate strategy of enabling anaerobic xylose fermentation, one that we predicted may augment stress tolerance. Perturbing sequence of the PKA regulatory subunit Bcy1 through simple protein fusion (with either a 260 amino acid auxin-inducible degron (AiD) tag without degradation capabilities or merely GFP) promotes anaerobic xylose fermentation equal to strain Y128 without the need for *HOG1* or *IRA2* deletion (93).

Since this strain retains functional Hog1 and grows well on glucose, we predicted the strain may have improved stress tolerance, which could enhance growth and xylose fermentation in 9% AFEX-pretreated corn stover hydrolysate (ACSH).

Here, we set out to dissect the contributions of regulators in the RAS/PKA and HOG pathways to anaerobic xylose fermentation, growth, and stress resistance. We demonstrate that while upregulating PKA is important for increased cell growth on glucose, *HOG1* deletion specifically benefits xylose utilization, in part by preventing phosphorylation of glycolytic enzymes by Hog1. Our results show perturbing Bcy1 sequence by protein fusion dramatically improved stress tolerance, as seen by increased growth in toxic 9% glucan-loading ACSH, but xylose fermentation remained blocked, suggesting physiological stress sensitivity is unlikely the cause of halted xylose consumption in concentrated hydrolysate.

## Methods

### Strains and Media

Strains used in this study are listed in Table 1. All strains expressed a codon-optimized cassette containing *XYLA* from *Clostridium phytofermentans, XYL3* from *Scheffersomyces stipitis*, and *TAL1* from *S. cerevisiae* (21). Generation of strains Y22-3, Y128, Y184, Y184 *ira2Δ*, Y184 *hog1Δ*, Y184 *ira2Δhog1Δ*, Y184 *bcy1Δ*, and 184 Bcy1-AiD was previously described (21,22,93). Y184 *ira2Δbcy1Δ* was made by replacing *BCY1* with *KanMX* marker in Y184 *ira2Δ*, and verified with diagnostic PCR*S. HOG1* was complemented in Y184 *ira2Δhog1Δ* o n a low-copy MoBY 1.0 plasmid (49). Site-directed mutagenesis was performed to express the *hog1^D144A^* or *hog1^A844Δ^* allele from the MoBY 1.0 plasmid, and correct mutants were verified by sequencing. Strains harboring *HOG1, hog1^D144A^*, or *hog1^A844Δ^* expressing plasmids or the empty-vector control were grown in media containing G418 to maintain the plasmid.

**Table 1.**
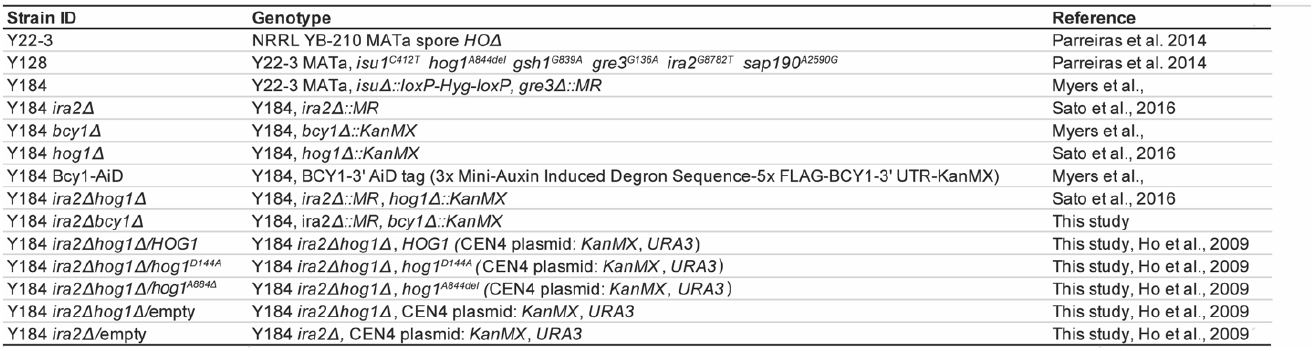
Strains used in this study.

YPDX medium was prepared as previously described (50) (1% yeast extract, 2% peptone, except that sugars were added at either 6% glucose and 3% xylose or 9% glucose and 4.5% xylose). Identified lignotoxins (51,52) were added to YPDX 6%/3% and the medium was sterilized by vacuum filtration. ACSH was prepared from *Zea mays* (Pioneer hybrid 36H56) grown in Field 570-C Arlington Research Station, University of Wisconsin and harvested in 2012, as previously described (53). Pretreated corn stover was hydrolyzed to either 6% or 9% glucan loading at 50°C for 5 days, and biomass was added 4 hours after hydrolysis began in 4 batches. The hydrolysate was centrifuged (2500xg for 30 minutes) and sterile filtered (0.22 μm pore size; Millipore Stericup). Final sugar concentrations were 53 g/L glucose and 21.7 g/L xylose for 6% glucan-loading ACSH, and 80 g/L glucose and 36 g/L xylose for 9% glucan-loading ACSH.

### TECAN Screening

Strains were grown to saturation in YPD 2% batch aerobically overnight, then diluted to an OD_600_ of 2 in YPD 2%. 5 μL of culture was added to 95 μL of 9% ACSH, YPDX 9%/4.5%, YPDX 6%/3%, or YPDX 6%/3% +LT in a Costar clear 96-well plate. OD at 600 nm was measured anaerobically for 48 hours in a TECAN Infinite 200 at 30°C, with measurements taken every 20 minutes (total of 144 measurement cycles) and multiple reads per well. Final OD was calculated based on the average OD reading per well per time point and using three biological replicates. Significant differences compared to Y184 were determined by performing a paired T-test at *p* < 0.05, pairing sample by replicate date.

### Batch culture fermentations

Overnight aerobic cultures grown in YPD 2% glucose were transferred to an anaerobic chamber, washed once with anaerobic medium, and inoculated into the tested media at an OD_600_ of 0.1. Fermentations occurred for 96 hours, with 1 mL aliquots removed throughout the time course for OD_600_ measurements and HPLC-RID analysis to measure glucose, xylose, and ethanol concentrations. Dry-cell weight biomass was measured by vacuum filtering cultures onto pre-weighed filters (0.45 μm pore size), microwaving on 10% power for 10 minutes, then drying in a desiccant for 24 hours before measuring. Biomass, OD, and concentrations of glucose, xylose, and ethanol were averaged from three biological replicates for each time point. ANOVA was used to determine significant differences in biomass, comparing 24h and 48h within each individual strain, at *p* < 0.05. Rates of sugar consumption were calculated by normalizing the change in sugar concentration to the fitted rate of biomass change during exponential (glucose) or stationary (xylose) phase, and a paired T-test was used to determine significant differences compared to Y184 at *p* < 0.05. Ethanol titer at 48h was averaged from three biological replicates.

### Phosphoproteomic analysis

Quantitative proteomic and phosphoproteomic samples of Y184 *ira2Δhog1Δ* and Y184 *hog1 A* were prepared as previously described using isobaric tandem mass tags (TMT) for phosphoproteomic analysis (93). Paired samples were collected from two independent replicates grown anaerobically in YPX 3% for three generations and harvested at OD_600_ ~0.5. Proteomic samples were analyzed by nanoflow liquid chromatography tandem mass spectrometry. COMPASS (54) was used to search against target-decoy yeast database (55). Raw data were transformed to log_2_ values, then the fold change between Y184 *ira2Δhog1Δ* and Y184 *ira2Δ* peptides was calculated. We focused on differentially abundant phosphopeptides, defined as those with a log_2_ difference of at least 1.5X in the same direction in both replicate comparisons. We used this threshold since TMT tagging is known to compress abundance differences (56).

## Results

Our goal was to dissect the contributions of different regulators in growth, xylose fermentation, and stress tolerance. We therefore measured fermentation and growth rates of a panel of strains grown in several concentrations of ACSH and laboratory media. To clarify the contribution of each regulator, we started with strain Y22-3, which overexpresses a codon-optimized cassette containing *XYLA* from *Clostridium phytofermentans, XYL3* from *Scheffersomyces stipitis*, and *TAL1* from *S. cerevisiae* but cannot utilize xylose, and strain Y184, which additionally lacks *ISU1* and *GRE3* and can thus respire xylose aerobically but not anaerobically (93). Both Y22-3 and Y184 can grow in the toxic 9% ACSH, whereas strain Y128, which can ferment xylose anaerobically, cannot (Fig.1). To test the impact of mutations that enable anaerobic xylose utilization, we generated Y184 derivatives lacking *IRA2, BCY1*, or *HOG1* individually as well as *IRA2 HOG1* or *IRA2 BCY1* in combination, to define how each mutation impacts stress tolerance, growth, and metabolism. We also included Y184 Bcy1-AiD, in which an auxin-inducible degradation sequence is fused to the C-terminal of Bcy1 (in the presence of functional *IRA2* and *HOG1*); it is important to note this sequence alone does not enable degradation of Bcy1 but imparts rapid xylose fermentation in lab medium without the need for *HOG1* deletion (93).

**Figure 1.**
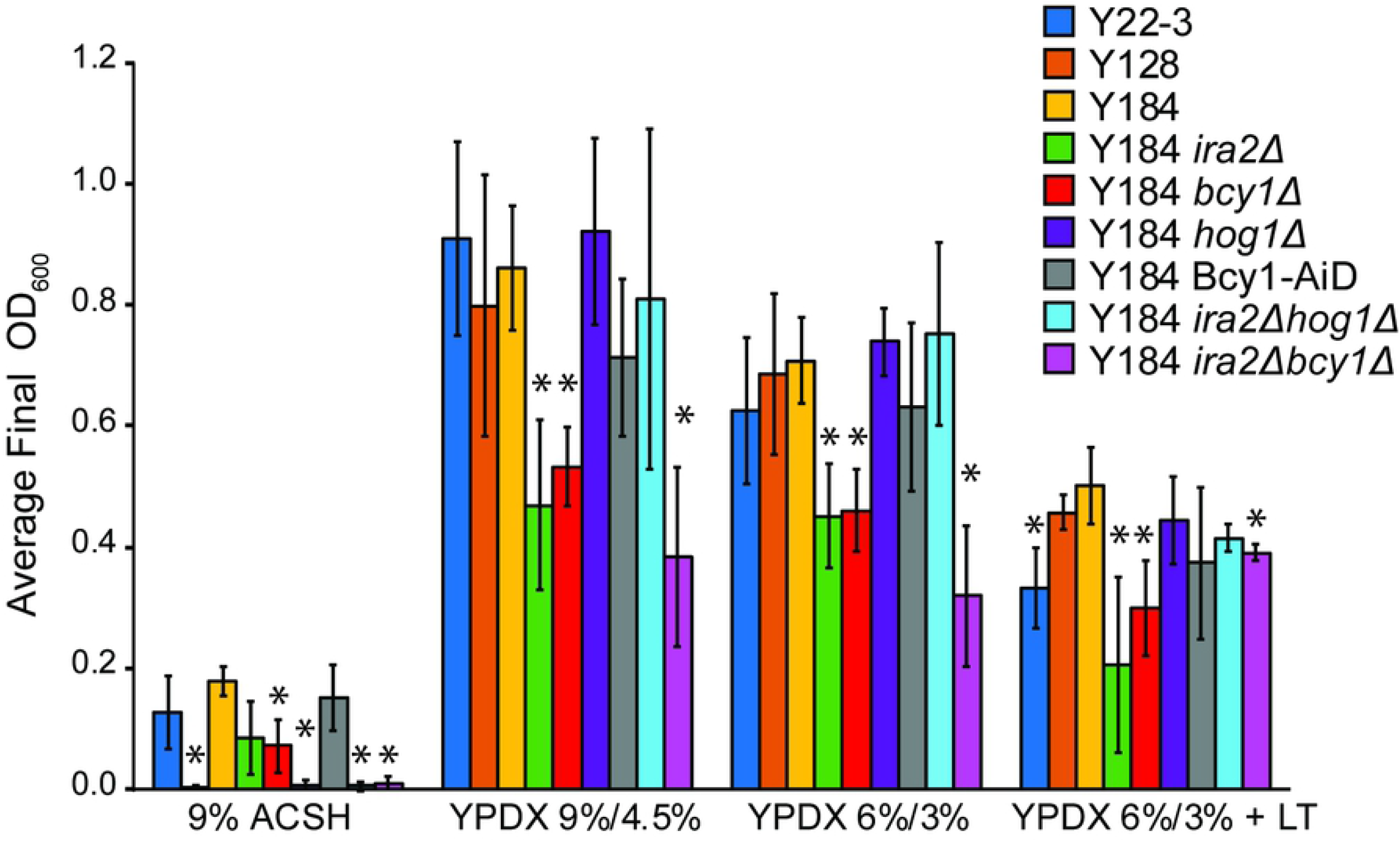
Stress susceptibility of strains engineered for xylose consumption. Strains were grown anaerobically in 9% ACSH, YPDX 9%/4.5%, YPDX 6%/3%, and YPDX 6%/3% + lignotoxins (LT) in 96-well plates, and final cell density at 48h was measured with a TECAN instrument. Average and standard deviations of final ODs were calculated from three biological replicates. Asterisks denote significant differences compared to Y184 grown in each respective condition (paired T-Test, *p* < 0.05).

### Improving stress tolerance does not restore xylose fermentation in toxic hydrolysate

We first measured tolerance of each strain to ACSH hydrolysates and rich laboratory medium containing mixed glucose/xylose with and without lignotoxins (LT), anaerobically in a 96-well plate reader. 9% ACSH is clearly inhibitory to growth, since all strains grew to much lower cell densities than in rich medium with lignotoxins added (Fig. 1). Parental strains Y22-3 and Y184 showed the best growth in 9% ACSH, as indicated by final cell density, even though they cannot use the xylose after glucose is consumed. Y184 *ira2Δ* a nd Y184 *bcy1Δ* s h owed reduced growth compared to Y184, but still grew; Y128, Y184 *hog1Δ*, Y184 *ira2Δhog1Δ*, and Y184 *ira2Δbcy1 Δ* were not able to grow. Thus, all strains lacking functional Hog1 showed increased sensitivity to 9% ACSH, as did the Y184 *ira2Δbcy1 Δ* strain. Since Hog1 functions in the osmotic stress response, we reasoned the high sugar content of 9% ACSH may be responsible for the sensitivity of *hog1Δ* strains. However, the mutants were not sensitive to rich medium with sugar concentrations matching 9% ACSH (9% glucose, 4.5% xylose, Fig. 1), indicating it is not the osmolarity of 9% ACSH that is inhibitory to these strains. Strikingly, Y184 cells with the Bcy1-AiD tag, which enables anaerobic xylose fermentation, displayed maximal growth in 9% ACSH, comparable to Y184. Therefore, as we predicted, Y184 Bcy1-AiD lacks the extreme stress sensitivity seen in *hog1Δ* strains. While adding lignotoxins to YPDX medium decreased growth, it was not as inhibitory to any strain as 9% ACSH, suggesting either the lignotoxin cocktail (52) added was lower than toxin levels in real hydrolysate or additional toxins not in our cocktail remain to be identified.

We next studied glucose and xylose consumption in 9% ACSH to characterize fermentation rates. We expected the strains whose growth was sensitive to the stresses of ACSH would ferment worse compared to tolerant strains. Surprisingly, we found all strains, even those containing functional Hog1, were incapable of fermenting xylose in hydrolysate generated at high-glucan loading (Fig. 2C, Fig. S1C). Y128 and Y184 *hog1 Δ* d i d not grow to densities as high as Y184, but Y184 Bcy1-AiD showed division as robust as Y184 (Fig. 2A), supporting observations seen with the 96-well plate screen. Consistent with our hypothesis, Y184 *hog1 Δ* a nd especially Y128 cultures showed reduced glucose consumption over the time course, whereas stress-tolerant Y184 and Y184 Bcy1-AiD depleted the glucose by 40 hours (Fig. 2B). Since we previously showed the Y184 Bcy1-AiD strain can ferment xylose anaerobically at higher rates than Y128 in laboratory medium (93) and has dramatically improved growth in 9% ACSH, we predicted Y184 Bcy1-AiD would display enhanced xylose consumption compared to Y128 and Y184 *hog1 Δ* i n 9% ACSH. Unexpectedly, this was not observed, as none of the strains used xylose in 9% ACSH (Fig. 2C) even though two fermented the glucose. Several strains marginally used xylose in 6% ACSH (Fig. S2C), but clearly ferment xylose in YPDX (see below), this suggests stresses in ACSH medium prevent xylose fermentation. Since the growth and glucose consumption of Y184 Bcy1-AiD is clearly recovered, these results suggest cellular stress is unlikely to be the cause of arrested xylose fermentation (see Discussion).

**Figure 2.**
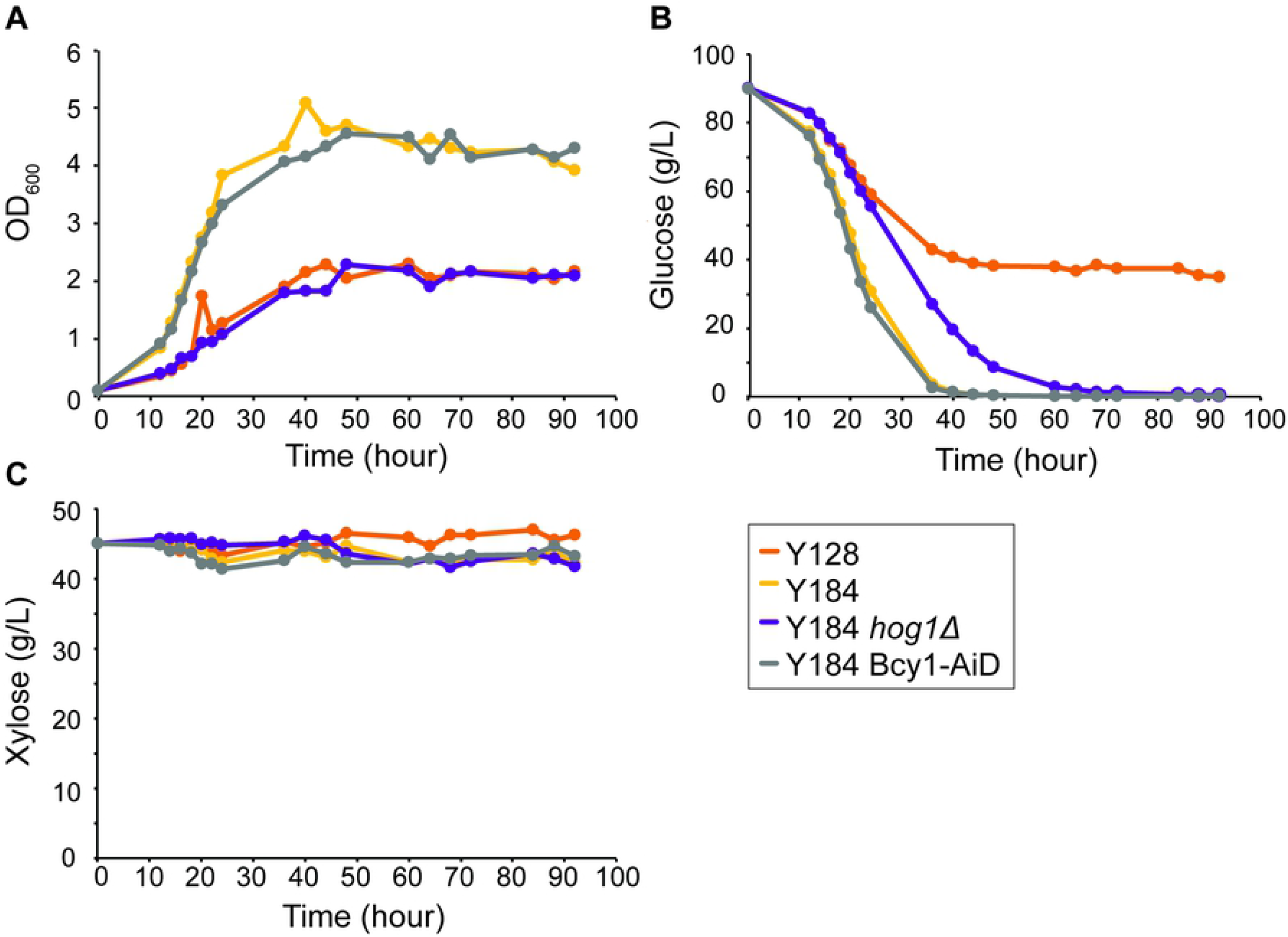
9% ACSH is inhibitory to xylose fermentation. Batch cultures of Y128, Y184, Y184 *hog1Δ*, and Y184 Bcy1-AiD were grown anaerobically for 92 hours in 9% ACSH. **A.** Average OD_600_ measurements from three biological replicates over time for denoted strains. **B.** Glucose concentration over time for strains grown in (A). **C.** Xylose concentration over time for the same cultures shown in (A).

### Differential contributions of regulators to glucose versus xylose fermentation rates

We next wanted to dissect the contribution of Ira2, Bcy1, and Hog1 to sugar fermentation. We worked with laboratory media supplemented with 60 g/L glucose and 30 g/L xylose to characterize fermentation when the strains are most productive, since they did not ferment xylose efficiently in ACSH. Interestingly, strains lacking Ira2 or Bcy1 or harboring the Bcy1-AiD fusion protein showed faster growth compared to the parental strains: all strains with these mutations reached maximum cell density by 24h, whereas Y22-3, Y184, and Y184 *hog1Δ* took 48h to reach maximal titers (Fig. 3A, Fig. S3A). Although it was not statistically significant in this growth assay, we noticed in the 96-well plate assay (Fig. 1) strains lacking *IRA2* or *BCY1* consistently grew to lower cell densities than the parental strains. Deletion of *HOG1* from Y184 reduced growth rate and glucose consumption rate compared to the other strains, whereas combined deletions of *HOG1* and *IRA2* produced a strain with fast growth rates and the ability to grow to a higher cell density than Y22-3 and Y184 (Fig. 3A,B, Fig. S3A,B). Thus, the beneficial effect of *IRA2* deletion overrides the deleterious effect of *HOG1* deletion for glucose-based growth.

**Figure 3.**
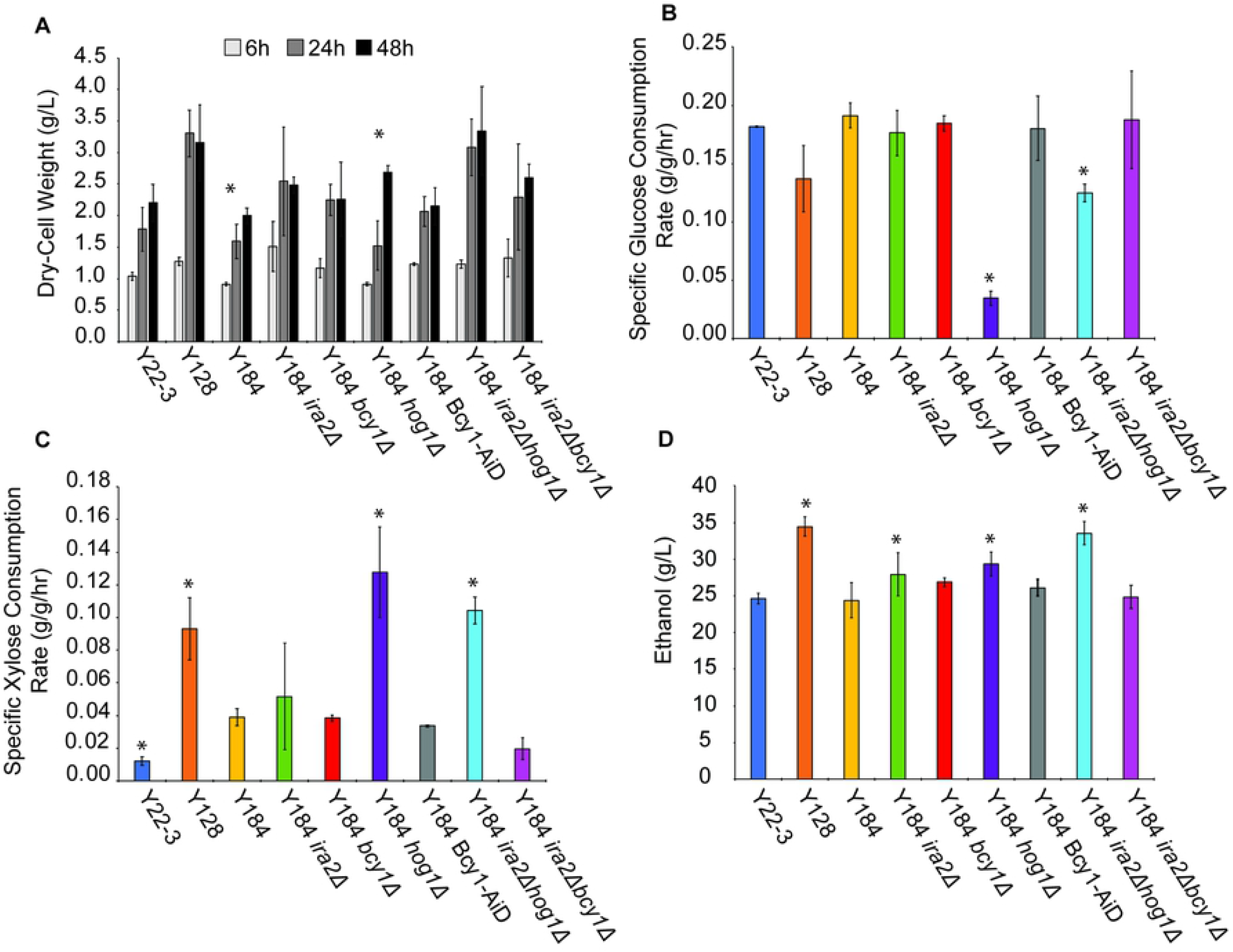
PKA activity increases growth, while *hog1Δ* improves xylose consumption. Batch cultures were grown in YPDX 6%/3% for 96 hours. Measurements are averages from three biological replicates. **A.** The average dry-cell weight biomass (g per L culture) measured at 6, 24, and 48 h post inoculation. Asterisk indicates a significant difference (p< 0.055) between the 24h and 48h timepoint within each strain. For all strains except Y184 and Y184 *hog1Δ*, the biomass accumulated was not significantly different at 48h compared to 24h, indicating faster saturation of those cultures. **B.** Specific glucose consumption rates calculated from the exponential phase of growth. **C.** Specific xylose consumption rate calculated from the stationary phase of growth. **D.** Ethanol titer at 48 hours post inoculation. For **B,C,D**, asterisks denote significant differences compared to Y184 (paired T-Test, *p* < 0.05).

While mutations in PKA regulators increase growth on and consumption of glucose, the opposite was observed for xylose consumption. Strains lacking *HOG1* had the highest rates of xylose consumption compared to the parental strains, whereas strains with PKA mutations alone did not have a significantly different xylose consumption rate from the Y184 parental strain (Fig. 3C). Thus, mutating *HOG1* is specifically important for efficient xylose fermentation. Moreover, Y128, Y184 *hog1Δ*, and Y184 *ira2Δhog1 Δ* b egin to ferment the xylose before glucose is completely depleted from the culture (Fig. S3B and C). Correspondingly, Y128, Y184 *hog1Δ*, and Y184 *ira2Δhog1Δ* had the highest ethanol titer at 48 hours (Fig. 3D, Fig. S3D). Y184 Bcy1-AiD was previously shown to use xylose at a comparable rate to Y128 in YPX medium when cultured at a high cell density (93). Interestingly, we discovered that the strain does not use xylose efficiently when the culture is started at low cell density (Fig. 3C), even though we recapitulate robust xylose utilization at high cell density in YPDX (Fig. S4A). The reason for this unique phenotype is not known; importantly, it did not explain the lack of xylose consumption in ACSH, since inoculating cultures at high cell titers did not improve xylose fermentation in the strain (Fig. S4B).

### Hog1 kinase activity decreases xylose fermentation

*HOG1* deletion allows for efficient xylose fermentation in YPDX. To test if it is the kinase activity of Hog1, as opposed to other effects of the Hog1 protein, we tested if the catalytically inactive *hog1-D144A* allele or truncated *HOG1* recapitulating Y128’s mutation *(hog1 A844Δ)* could block xylose fermentation when introduced into Y184 *ira2Δhog1Δ*, equal to reintroducing the functional *HOG1* allele. We saw the catalytic activity of Hog1 was required to suppress anaerobic xylose fermentation: reintroducing wildtype *HOG1* increased glucose consumption and inhibited xylose fermentation, whereas expression of catalytically inactive or truncated Hog1 did not significantly affect the glucose or xylose consumption rate, since the strains matched the performance of Y184 *ira2Δhog1Δ* (Fig. 4). This reveals Hog1 kinase activity normally inhibits xylose fermentation, suggesting Hog1 actively phosphorylates one or more targets to prevent xylose utilization.

**Figure 4.**
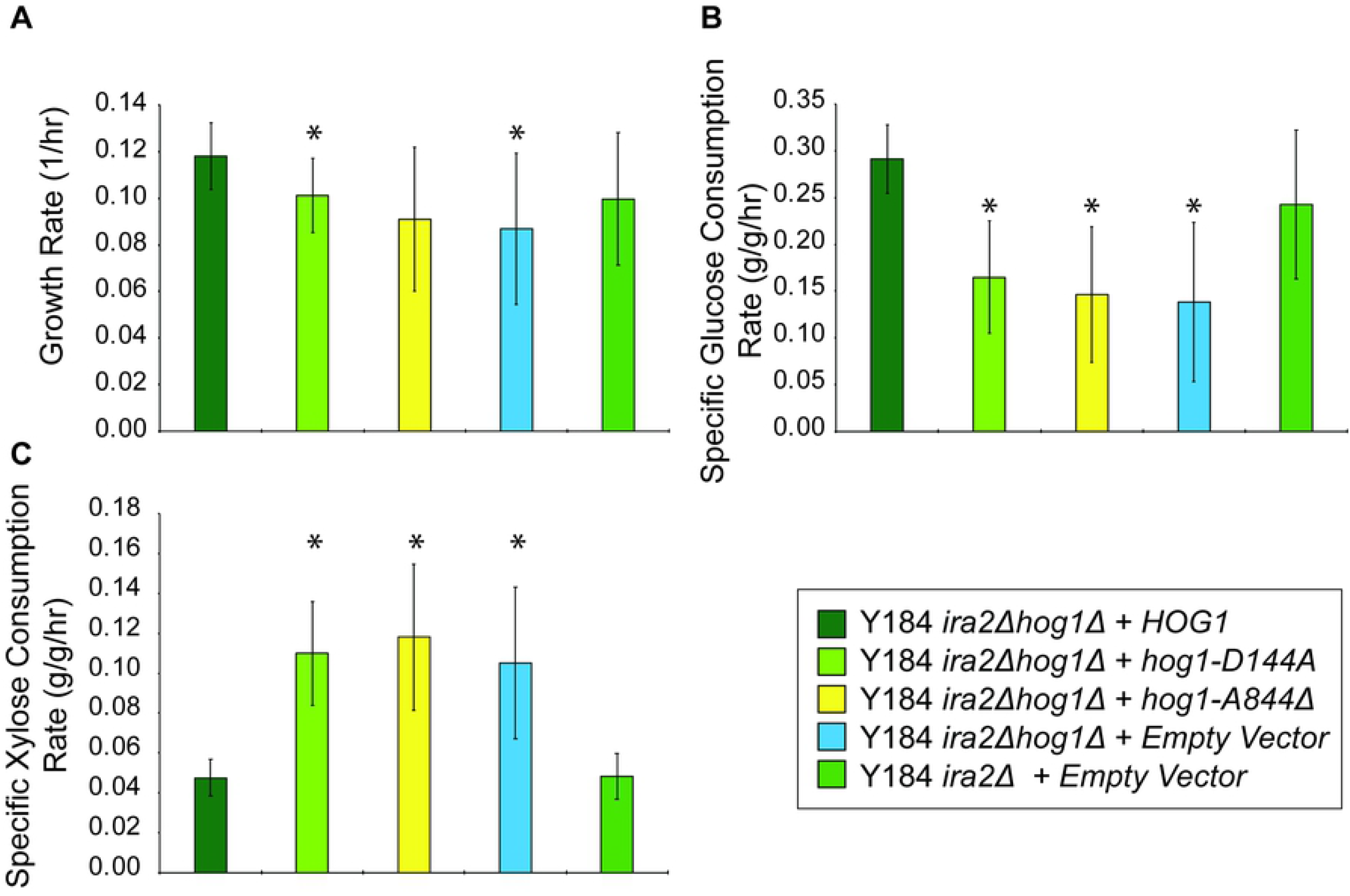
Hog1 activity blocks xylose consumption. Wildtype, catalytically inactive (*hog1^D144A^*), and truncated (*hog1^A844Δ^*) *HOG1* alleles were expressed in Y184 *ira2Δhog1 Δ* cells and grown in batch culture for 96 hours in YPDX 6%/3%. Rate measurements were based on average and standard deviation from three biological replicates. **A.** Average growth rate calculated from the exponential phase of growth. **B.** Specific glucose consumption rate calculated from the exponential phase of growth. **C.** Specific xylose consumption rate calculated from the stationary phase of growth. Asterisks denote significant differences from Y184 *ira2Δhog1Δ/HOG1* (paired T-Test, *p* < 0.05).

We performed phosphoproteomic analysis to implicate potential Hog1 targets impacting xylose consumption. Y184 *ira2Δ* and Y184 *ira2Δhog1 Δ* were grown anaerobically in batch culture in YPX medium, and phospho-proteomes were measured by quantitative mass-spectrometry (see Methods). We identified only 11 phosphopeptides whose abundance was reproducibly lower in Y184 *ira2Δhog1Δ* (Table 2, see methods). Of these 11, four are linked to metabolism: E1 alpha subunit of pyruvate dehydrogenase (Pda1), pyruvate kinase (Cdc19), glyceraldehyde-3-phosphate dehydrogenase (Tdh1), and trehalose-6-phosphatase complex (Tsl1). Pda1, Cdc19, and Tdh1 have roles in glycolysis, whereas Tsl1 generates the overflow metabolite trehalose. In contrast, eight phosphopeptides showed reproducibly increased abundance in Y184 *ira2Δhog1Δ*. Of these eight, four are from proteins with unknown functions, one is a heat shock protein (Hsp26), two are components of the 40S ribosome (Rps6A, Rps6B), and one is an epsin-like protein involved in endocytosis (Table 2). Several of these enzymes, including Cdc19 and Tsl1, directly interact with Hog1, and others harbor the known Hog1 consensus sites around the affected residue, suggesting Hog1 directly phosphorylates these proteins (see Discussion).

**Table 2:** Peptides with phosphorylation changes in Y184 *ira2Δhog1Δ*

| Protein | Common Name | Phospho Site | Direction of change in Y184 *ira2 Δhog1 Δ* |
| --- | --- | --- | --- |
| YJL136C | RPS21B | S65 | Decreased |
| YER178W | PDA1 | S336 | Decreased |
| YER178W | PDA1 | S344 | Decreased |
| YDL223C | HBT1 | S41 | Decreased |
| YMR031C | EIS1 | S736 | Decreased |
| YAL038W | CDC19 | S22 | Decreased |
| YMR031C | EIS1 | T767 | Decreased |
| YDL223C | HBT1 | S363 | Decreased |
| YJL052W | TDH1 | T136 | Decreased |
| YPL118W | MRP51 | S327 | Decreased |
| YML100W | TSL1 | S266 | Decreased |
| YBR072W | HSP26 | S208 | Increased |
| YNL115C | YNL115C | S244 | Increased |
| YNL115C | YNL115C | S42 | Increased |
| YDL161W | ENT1 | S317 | Increased |
| YBR181C | RPS6B | S232 | Increased |
| YPL090C | RPS6A | S232 | Increased |
| YLR413W | INA1 | S654; S657 | Increased |
| YLR413W | INA1 | S652; S657 | Increased |

## Discussion

Our results provide new insights into how mutation of different signaling proteins can impart distinct physiological responses that, when integrated, improve anaerobic conversion of sugars to products, in this case ethanol. We show mutations in RAS/PKA signaling that up-regulate PKA activity enhance growth and perhaps glucose consumption rates, but possibly at a cost of final cell density, whereas deletion of HOG1 is essential for xylose fermentation but at the expense of hydrolysate tolerance. Together, these results suggest several important results relevant to lignocellulosic fermentation.

PKA and Hog1 contribute separable features to xylose fermentation. Increased PKA activity is known to increase cell growth rate and glucose consumption (57). PKA upregulates expression of ribosome biogenesis genes, which supports rapid growth (58,59). Moreover, it promotes glycolytic flux by inducing transcription of genes and phosphorylating enzymes involved in glycolysis (60–62). We recently showed PKA also has roles in the hypoxic response when cells are grown anaerobically on xylose (93). This response is mediated by the Azf1 and Mga2 transcription factors and may further promote glycolytic flux. Therefore, it is consistent that we find strains with deletions of RAS/PKA regulators display faster growth (Fig. 3A, Fig. S3A). Despite the increased growth and glucose consumption rates when PKA regulators are deleted, this effect is not as dramatic with xylose. While strains with mutations in *IRA2* or *BCY1* do consume more xylose than Y184 (Fig. S3C), the rates of consumption are not significantly faster than Y184 (Fig. 3C). The reduced final-cell titers seen in strains with up-regulated RAS/PKA may result from an inability to accumulate storage carbohydrates. During the diauxic shift, glycogen storage occurs when glucose has been depleted to fifty percent of its starting concentration (63). However, increased PKA activity prevents accumulation of storage carbohydrates, such as glycogen and trehalose, by inhibiting expression of biosynthesis enzymes and activating catabolic enzymes (64–68). In our strains lacking RAS/PKA inhibitors, it is possible upon glucose depletion, the cells lack stored carbohydrates to metabolize, limiting cell titers compared to strains with functional *IRA2* and *BCY1* (Fig. 1, Fig. S3A).

In contrast to PKA upregulation, *HOG1* deletion specifically affects xylose utilization. Our results show that under standard conditions, even in the absence of added stress, Hog1 kinase activity inhibits xylose utilization, perhaps by directly phosphorylating glycolytic enzymes (Fig. 4C, Table 2). Although typically thought of as a stress regulator, Hog1 has recently been implicated in the response to glucose (28,29,69) and was shown to phosphorylate the glycolytic enzyme Pfk26 (70) and physically interact with Cdc19, Pfk1, and Pfk2 (71). Hog1 also plays an important role in glycerol production, which may influence glycolytic flux to steer production from pyruvate towards glycerol (72–77). The enzymes we detected with significant phosphorylation differences upon *HOG1* deletion function lower in glycolysis, after the entry point of xylose via the pentose phosphate pathway (Table 2), but we cannot exclude that other enzymes or regulators are also affected. Modeling of glycolytic flux during osmoadaption suggests Hog1 activity stabilizes pyruvate production to prevent starvation (75), which may account for the decreased phosphorylation of Cdc19’s activation site in *hog1Δ* mutants (Table 2).

An interesting question is the relationship between PKA and Hog1 activity. We previously found Y184 *bcy1 Δ cells* display reduced phosphorylation of the Hog1 protein on its activating site when grown anaerobically in YPX medium (93), suggesting a PKA-dependent mechanism of inhibiting Hog1 activity to allow xylose fermentation. Hog1 was shown to be activated during glucose depletion (28,69). It is possible as cells begin to deplete glucose from YPDX, Hog1 becomes activated in Y184 but that this is suppressed when PKA is up-regulated via *BCY1* or *IRA2* deletion. The largest effects on xylose metabolism occur when PKA upregulation and *HOG1* deletion are combined. Unfortunately, a byproduct of this is likely causing extreme stress sensitivity. Strains lacking *HOG1* (Y128, Y184 *hog1Δ*, and Y184 *ira2Δhog1Δ)* are especially sensitive to hydrolysate (Fig. 1, Fig. 2A, Fig. S1A). Moreover, the combination of upregulated PKA and *HOG1* deletion may exacerbate stress sensitivity, since PKA suppresses the stress response (24), while Hog1 activates it (25). These strains are able to ferment xylose in favorable conditions (Fig. 3C, Fig. S3C), but are unable to ferment the sugar in toxic 9% ACSH (Fig. 2C, Fig. S1C).

We hypothesized improving stress tolerance would improve xylose fermentation in stressful conditions. We predicted Y184 Bcy1-AiD, which harbors functional *HOG1* but can ferment xylose anaerobically in rich medium when started at high titers (93), would display both higher stress tolerance and anaerobic xylose fermentation in hydrolysate. Interestingly, while Y184 Bcy1-AiD clearly grew well in 9% ACSH, it did not utilize the xylose even when started at high cell titer (Fig. 1, Fig. 2A, Fig. 2C Fig. S4B). This suggests stress sensitivity is not the sole factor limiting xylose fermentation. There are many types of lignotoxins present in hydrolysates (51), and while their effects are somewhat understood, many of their specific targets remain to be elucidated (33). One possibility is that toxins are directly inhibiting enzymes required for anaerobic xylose fermentation, as shown in *Escherichia coli* where feruloyl and coumaroyl amides were discovered to be allosteric inhibitors of *de novo* nucleotide biosynthetic enzymes (78). Jayakody et al. (2018) found that glycoaldehyde and methyglyoxal present in hydrolysate are key inhibitors of xylose fermentation (79). Furthermore, acetic acid is known to decrease enolase activity, ultimately slowing down glycolysis (80). This has led to other groups engineering yeast to reduce and ferment acetate or increase acetate tolerance (43,81,82). More studies identifying the targets of lignotoxins will help to clarify the bottleneck in xylose metabolism when microorganisms are grown in hydrolysate media.

Our results also revealed unexpected information on the different routes of PKA regulation with regard to the stress response. Removing either *IRA2* or *BCY1* had mild impacts on ACSH tolerance, but deletion of both genes greatly impacted tolerance of 9% ACSH (Fig. 1, Fig. 2A, Fig. S1A). This suggests there are multiple lines of partially redundant regulation of stress tolerance by the RAS/PKA pathways. Upregulation RAS by *IRA2* deletion is predicted to increase cAMP and thus PKA activity (83), whereas deletion of *BCY1* removes the cAMP-responsive inhibitor of PKA. One possibility is PKA is only partially upregulated by single-gene deletion, but double-deletion of *IRA2* and *BCY1* produces much stronger activation and thus complete suppression of the stress response. There are other methods of RAS/PKA pathway regulation which support the possibility that single deletion of *IRA2* or *BCY1* causes only a partial upregulation. The RAS/PKA pathway undergoes feedback inhibition to control cAMP concentrations through predicted PKA-directed phosphorylation of Cdc25 and Pde1 (84). Furthermore, the interaction between the catalytic and regulatory subunits of PKA is regulated by other factors, including kelch-repeat proteins Krh1/2 (85). Another possibility is PKA may be directed to different substrates in different situations. Beyond allosteric cAMP regulation influenced by RAS, Bcy1 is also regulated by phosphorylation, which can influence PKA substrate specificity and localization (86–90). Recent studies in mammalian cells show the negative regulator of PKA does not dissociate from the active kinase at physiological cAMP levels (91), and the substrate determines the dissociation rate of catalytic and regulatory subunits (92). Thus, activation of PKA via Bcy1-cAMP binding may provide different effects than if Bcy1 is missing from the cell. Finally, it is also possible that PKA-independent effects of *IRA2* deletion separately regulate the stress response. Future studies to dissect these and other effects will contribute to our understanding of how to engineer cells for anaerobic xylose fermentation in lignocellulosic hydrolysates.

## Acknowledgments

We thank Trey Sato for providing strains, Mick McGee for HPLC analysis, and members of the Gasch lab for constructive discussions.

## Author Contributions

Conceptualization: ERW, KSM, and APG. Investigation: ERW, KSM, and NMR. Methodology: ERW, KSM, NMR, JJC, and APG. Formal Analysis: ERW, KSM, and APG. Funding Acquisition: APG and JJC. Project Administration: APG. Resources: APG and JJC. Software: KSM, NMR, JJC, and APG. Supervision: APG. Validation. ERW, KSM, and APG. Visualization: ERW, KSM, and APG. Writing-Original Draft Preparation: ERW and APG. Writing - Review & Editing: ERW, KSM, and APG.

## Supplementary Figure Legends

**Supplementary Figure 1. 9% Growth and metabolism profiles in 9% ACSH.** Batch cultures were grown in 9% ACSH anaerobically for 92 hours. Data represent average and standard deviation from three biological replicates. **A.** OD_600_ measurements over time. Glucose (**B.**) and xylose (**C.**) concentration in the media over time.

**Supplementary Figure 2. Growth and metabolism profiles in 6% ACSH.** As described in Supplementary Figure 1 except for anaerobic 6% ACSH growth, measuring dry-cell weight (**A.**), and glucose (**B.**), xylose (**C.**), and ethanol (**D.**) media concentration over time.

**Supplementary Figure 3. Growth and metabolism profiles in YPDX 6%/3%.** As described in Supplementary Figure 1 except for anaerobic YPDX 6%/3% growth, measuring dry-cell weight (**A.**), and glucose (**B.**), xylose (**C.**), and ethanol (**D.**) media concentration over time.

**Supplementary Figure 4. High starting cell titers increases xylose consumption in nutrient-rich medium, but not ACSH.** Batch cultures were grown anaerobically for 96 hours in YPDX 6%/3% (**A.**) or 6% ACSH (**B.**). Cultures were started at an OD_600_ of 3. Data represent average and standard deviation of three biological replicates. Comparing Panel A to Figure 3C shows that the Y184 Bcy1-AiD strain ferments xylose when the culture is inoculated at a higher starting OD but not when inoculated at a lower cell density.

